# Improving Children’s and Their Visitors’ Hand Hygiene Compliance

**DOI:** 10.1101/355818

**Authors:** D. Lary, A. Calvert, B. Nerlich, J. Segal, N. Vaughan, J. Randle, K. R. Hardie

## Abstract

**Background:** Numerous interventions have tried to improve healthcare workers’ hand hygiene compliance, however little attention has been paid to children’s and their visitors’ compliance.

**Aim:** To increase children’s and visitors’ compliance using interactive educational interventions.

**Methods:** This was an observational study of hand hygiene compliance before and after the introduction of educational interventions. Qualitative data in the form of Questionnaires and interviews was obtained.

**Findings:** Hand hygiene compliance increased by 21.4% (P <0.001) following the educational interventions, with children’s compliance reaching 40.8% and visitors’ being 50.8%. Compliance varied depending on which of the five moments of hygiene was observed (P<0.001), with the highest compliance after body fluid exposure (96%). Responses from questionnaires showed educational interventions raised awareness of the importance of hand hygiene (69%, 57%) compared to those who had not experienced the educational intervention (50%).

**Conclusion:** Educational interventions may result in a significant increase in children’s and visitors’ hand hygiene (P <0.001).

## Background

Children are vulnerable to infectious diseases (Willmott et al., 2016) and NICE 2017 (NICE, 2017) calls for education providers and parents to do more to promote good hand hygiene practices. This is especially relevant when considering children’s vulnerability in healthcare settings where not only are children treated by a plethora of healthcare workers who travel in and out of different clinical settings, but they are typically surrounded by ill people. Consequently the healthcare environment has been emphasised as a potential source of harm for patients in the last few years and the reduction of healthcare associated infections (HCAI) is now part of the everyday delivery of healthcare treatment.

To prevent and reduce HCAI transmission, it is important to determine if the main routes of exposure to infection are direct, indirect, or due to repeated person-to-person contact. In children, the transmission of infections is likely to correlate with their natural behaviour (e.g. regular exploration of their mouths). The resultant spread of respiratory secretions coupled with an immature immune system combine to increase children’s risk of infections (Snow et al., 2008) and they are especially at high risk of respiratory infections and gastrointestinal diseases (Stein et al., 2007).

Hand hygiene (HH) is the single most important measure for reducing HCAI, and interventions can improve compliance (Randle et al., 2010) with the most effective being multimodal. (Naikoba and Hayward, 2001; Gould et al., 2017)

Unsurprisingly studies have focused on Healthcare workers’ compliance and patients’ and visitors’ has been overlooked, even though their Hand Hygiene Compliance (HHC) is important, especially if they augment the care provided by the HCWs as a parent would. Patients and visitors pose a high risk because of their potential to (i) transmit virulent pathogens from the community to the healthcare setting and/or (ii) transfer pathogens within clinical areas to the patient (directly or indirectly). (Gould et al., 2017; Randle et al., 2010; Munoz-Price et al., 2012)

This study monitored children’s and their visitors HHC before and after the introduction of an educational intervention (Supplementary Figure A) The educational intervention was either the Glo-yo (Supplementary Figure B, Supplementary Figure C, Supplementary Table A); or a video.

## Methods

### Ethical and Regulatory Aspects

The Research Ethical Committee (REC) Committee East Midlands Research NHS and the Research & Innovation department, NHS, approved this study.

### Study design

This observational study was conducted on six paediatric wards in a teaching Hospital in the East Midlands. Random sampling (slips of paper in a hat) allocated two paediatric wards for each educational intervention (the Glo-yo or the video) and the control group which received no intervention (see Supplementary Table A). The baseline phase included HHC rates using the WHO 5 moments of hand hygiene (2009). The intervention phase included Hand HHC rates and the educational interventions. After the interventions, a qualitative questionnaire was given to the parents/carers of the children (3-15 years) or children (≥16). Questions asked about HH behaviours, beliefs and attitudes about infection, hygiene and cleanliness that may influence or prevent effective HH, and views about different HH approaches, including the use of the Glo-yo or Video.

### Statistical analysis

The data were analysed using SPSS statistic software (IBM SPSS statistic v. 21) and GraphPad Prism6. HHC rates composed of simple frequency counts and Chi-square tests. The questionnaire responses were collated in categories inherent in the questions themselves, compared using simple frequency counts and grouped into themes.

## Results

### Baseline

A total of 525 HH opportunities of patients and visitors were monitored, and the overall compliance rate was 157/525 (30%, Table IA: proportion complied). HHC was low, particularly for children (10%). This rate was significantly different from that of their visitors (26%: P< 0.05). There was also a significant difference in HHC dependent on the moment of HH, irrespective of whether they were children or visitors (P< 0.001). The lowest level of compliance was observed after contact with patient surroundings (13%), and the highest was after exposure to body fluid (100%). Similarly, HHC of patients and visitors depended on the ward that they were on (P = 0.31) and were significantly different dependent on the time of day (P <0.001).

**Table 1.**
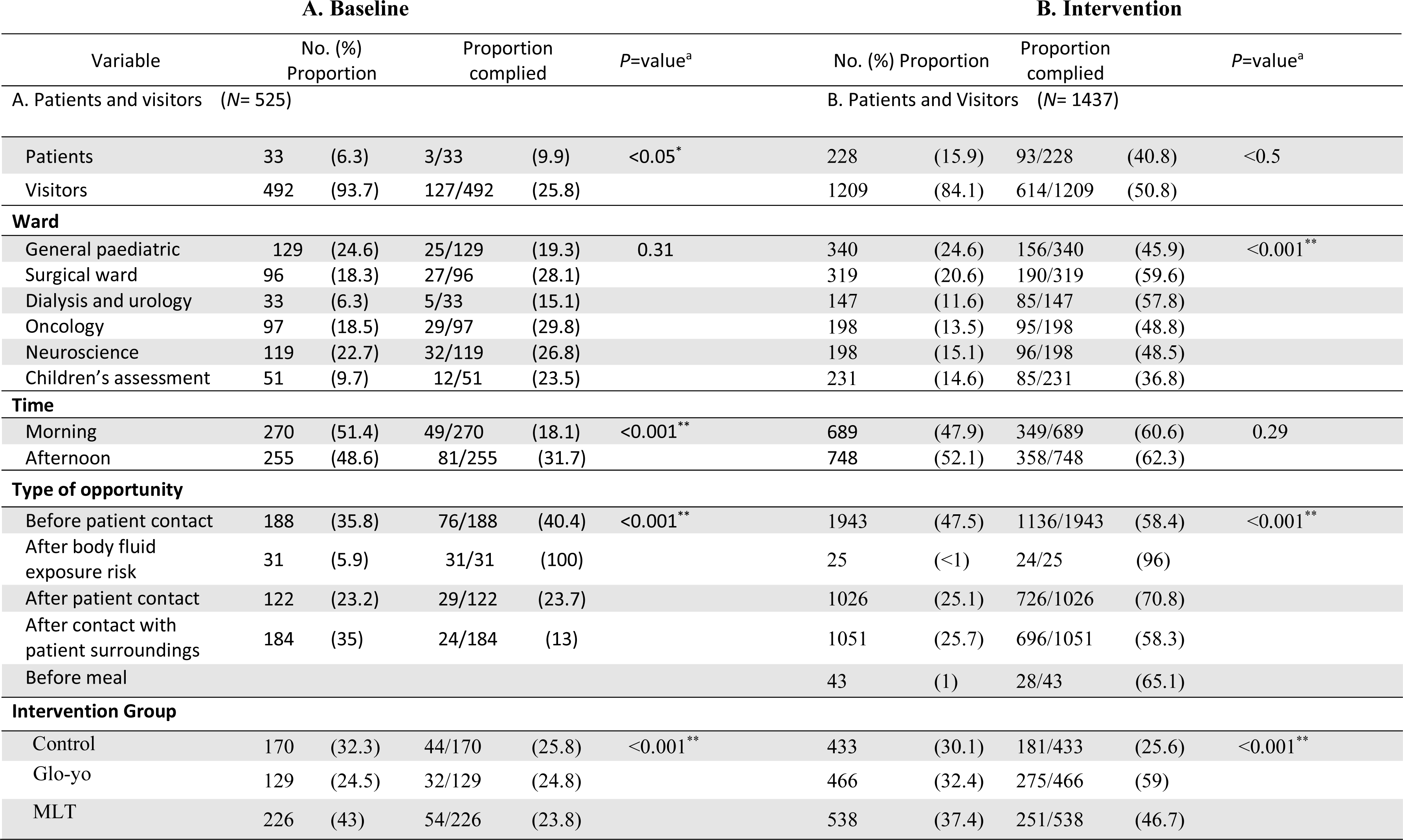
A X^2^-Test of difference in proportions of opportunities adhered to, across levels of variable. Left Column (A) shows the baseline data and right Column (B) the intervention ( intermidate phase data) *significant (P < 0.05) **highly significant (P < 0.001)

**Table 2.** Participant comments in response to two of the questions on the questionnaire.

| Intervention group | Comments on the question | Comments on the question |
| --- | --- | --- |
|  | 'How would they describe the session and the activities carried out by the facilitator?' | 'Would you recommend the sessions to children in schools' |
| Glo-yo | *'The session is very effective though the time is not enough to explore the new toy (Glo-yo).'* | *'Strongly agree, schools will be much better to learn about hand washing'* |
|  | *'I found the session very interesting and very encouraging and very easy task to do'.* | *'Very interesting would like to have same session in schools'* |
|  | *'Very helpful, the toy was interesting to use and good way to teach kids hand washing.'* | *'Mum found it interesting and very worthwhile. I would recommend it to school children. My daughter found it interesting'.* |
|  |  | *'I would recommend the hand wash in schools it will encourage children more to wash hands and be germ free than hospital'.* |
| MLT | *'The session in general is encouraging for children to learn about hand washing and germs. The use of phone is not easy for my child to understand'* | *'For my child was too much information, i enjoyed the video as a parent'.* |
|  | *'The video is too long and not easy to follow by child'* | *'The session was encouraging!'* |
|  | *'The video is very complicated for a child to understand. I enjoyed the video as it's easier to follow and understand'.* | *'The session with the facilitator was fun hand washing session was useful. However the phone video is not easy for child to understand'.* |
| Control |  | *'Session was helpful in understanding when to wash hands, and how, will recommend for school children'.* |
|  |  | *'Helpful session should be more practical to apply in schools'.* |

### Post intervention phase

1437 HH opportunities were observed. HHC increased by 24.4% compared to the baseline phase, and was significantly different between (i) children and visitors (*P*<0.01), (ii) the moments of contact providing the opportunity, (iii) the type of paediatric ward observed, and (iv) the intervention used (*P*< 0.001) (Table IB). The higher HHC in the afternoon shift was not significantly different from the morning shift (P = 0.29). HHC of patients and visitors in both intervention groups (but not the control group) was significantly different to the baseline phase HHC (*P*<0.001). The control group had similar HHC during the intervention phase (30.1%) compared to the baseline (32.3%). Interestingly HHC improvement was greatest after the intervention session using the Glo-yo, and this was a statistically significantly difference (*P*<0.001).

### The intervention session was successful at raising awareness of the importance of Handwashing

Of the 62 children and visitors approached, 31 agreed to participate in the educational intervention. The Glo-yo group included 16/31 (51.6%) of the participants (9/16 were patients). The Video group included 7/31 (22.5%) of the participants (5/7 patients). The control group included 8/31 (25.8%) of the participants (1/8 patients) (who only had access to HHC leaflets). All children were given a questionnaire to complete to determine their perception of the intervention session.

Children reported that the educational interventions raised their awareness of hand hygiene, with the Glo-yo intervention prompting a higher proportion of the participants to indicate that they strongly agreed with this (Figure 1).

**Figure 1.**
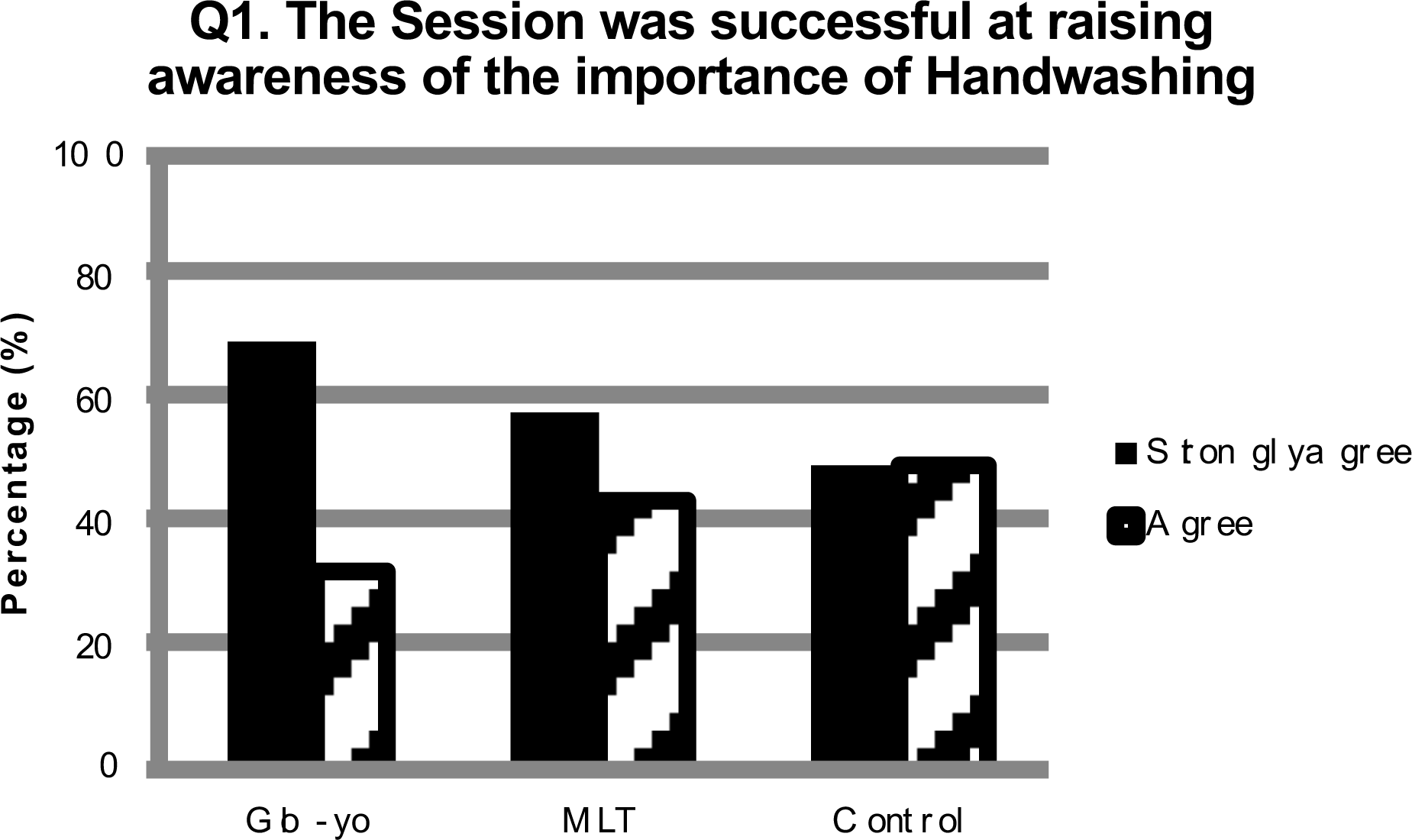
Participant feedback indicated that interactive sessions raised their awareness of the importance of Handwashing.

### The intervention session helped increase children’s knowledge and understanding of germs and handwashing

The questionnaire sought participant feedback on; A. why we wash our hands, B. germs and bacteria, C. when to wash hands, and D. parts of hands that are difficult to wash. The answers varied between intervention and subcategory of question. The Glo-yo intervention group agreed strongly with respect to all question subcategories (Figure 2).

**Figure 2.**
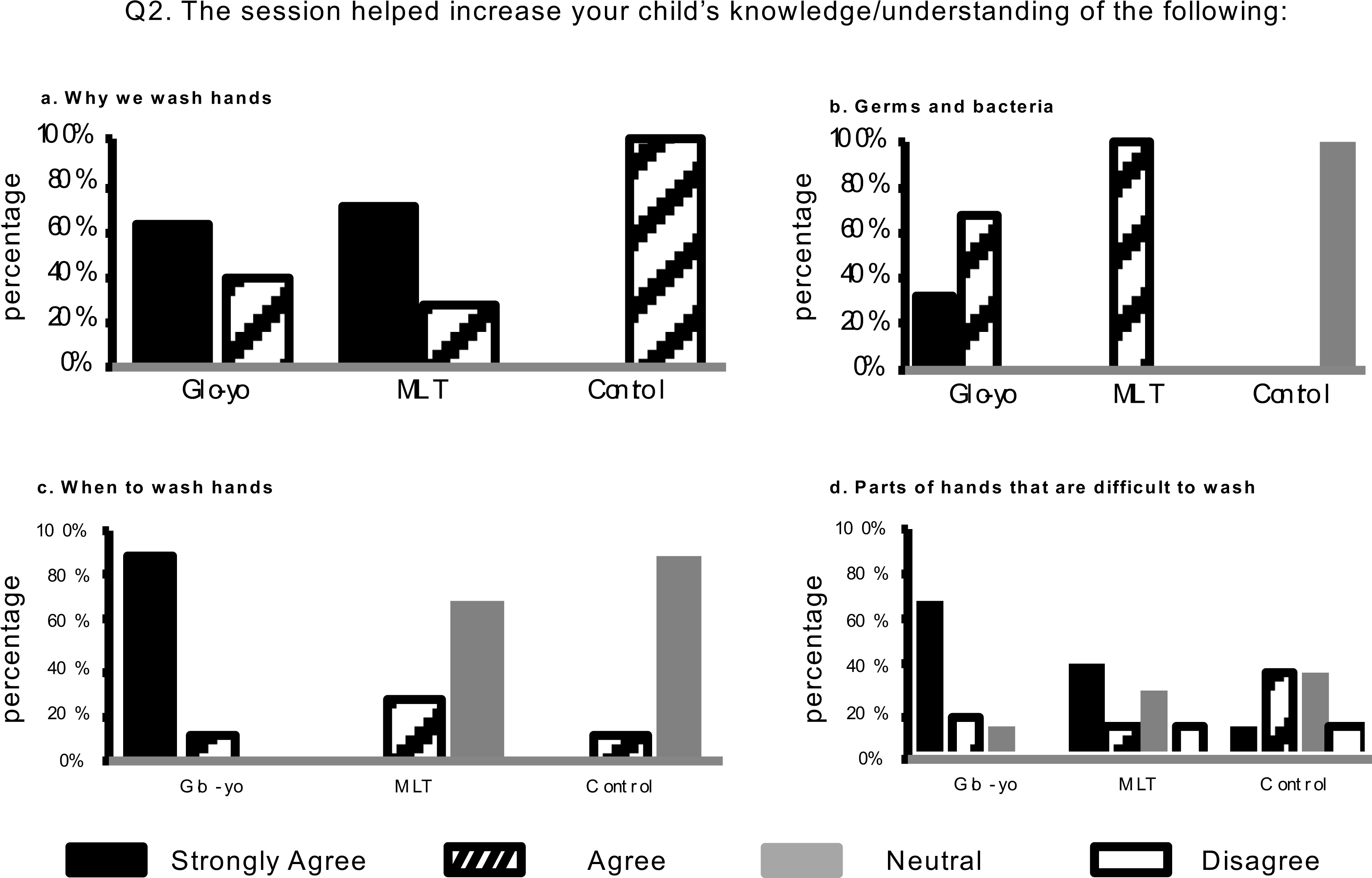
Participants agreed that the intervention sessions helped increase children’s knowledge and understanding of germs and handwashing. The responses to the second question on the questionnaire are shown (‘The session helped increase your child’s knowledge/understanding of the following:’).

Almost two thirds of participants in the Glo-yo and MLT intervention groups strongly agreed that the session and both training aids focused on why we wash our hands (62.5% and 71.4%), but 100% of the control group merely agreed with this (Figure 2a). When asked about whether the intervention increased knowledge about bacteria and germs, 33.3% of the participants in the Glo-yo group highly agreed and 100% of the Video group agreed, which contrasted with the control group, who were 100% neutral on this point (Figure 2a). When the participants considered whether the intervention sessions dealt with when to wash hands, 88% of the Glo-yo group strongly agreed, whereas 71% of the Video group and 88% of control group were neutral (Figure 2c). Finally, when asked whether the intervention session increased the knowledge and understanding of the parts of hands that are difficult to wash, 69% of the Glo-yo group, 43% of the Video group and only 13% of the control group strongly agreed. Indeed, a small proportion of the participants of the Video and controls disagreed with this (Figure 2d).

### The session improved children’s handwashing even for one day

Due to the limited time that patients spend in hospital, and because the session was only performed once with each participant, the final part of the questionnaire aimed to determine whether a single intervention session could improve handwashing even for one day. More than half of the Glo-yo group strongly agreed 56% whilst the participants of the other intervention groups were less convinced (Figure 3).

**Figure 3.**
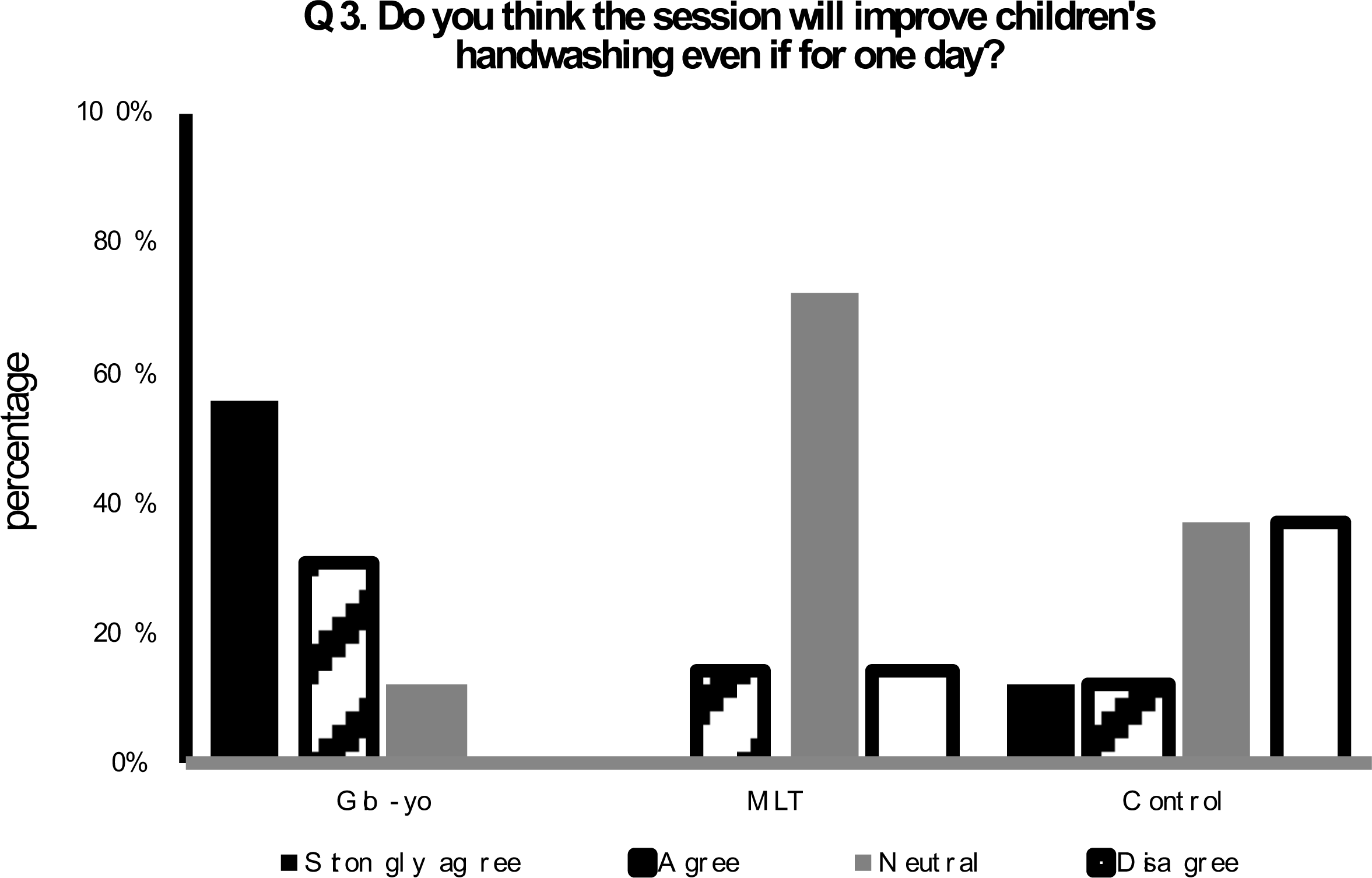
Participants agreed that the intervention sessions would be effective in improving children’s handwashing even for one day, with the strongest positive response being for the Glo-yo intervention.

## Discussion

Studies focusing on the HHC of patients and visitors in healthcare settings are limited (Buet et al., 2013). However, previous studies have reported an increase in HHC after educational intervention (McGuckin et al., 1999; Chen and Chiang, 2007; Fishbein et al., 2011).

Children and visitors had the highest level of compliance ‘after exposure to their own body fluids which has previously been identified (Randle et al., 2010) This may be as a result of self-protection, or due to emotional sensations including feelings of unpleasantness, discomfort and/or disgust (Whitby et al., 2007). The lowest compliance was found for the moment ‘after contact with patient surroundings’. This increased after intervention by 45% to reach 58.3%. Although this is considered a low compliance rate, it is significantly higher than recent data (Randle et al., 2013), and it is important as near touch sites pose the highest risk to patients, especially those in close and direct contact with patients (Dancer, 2009). Another high increase in HHC was observed ‘after contact with patients’. This was mainly observed in visitors, increasing from 23.7% to 70.8%, to reach a level >20% higher than previous observational studies (Randle et al., 2010). No study was found that looked at HHC of patients before a meal, in this study it was observed that compliance at this opportunity at the intervention phase was as high as 65 %.

This study indicates that HHC is better than previously reported, and provides evidence of a significant increase in HHC during intervention (P <0.001). The Glo-yo session proved the most successful intervention and was able to raise awareness of the importance of HH, with parents strongly agreeing that the Glo-yo session will improve their child’s hand washing. This aligns with previous research indicating educational and psychological programmes integrating tangible materials and images of the subject to be learnt can improve motivation and learning with the added benefit of long term behavioural change (Bairaktarova et al., 2011; Worthington et al., 2001; Ho et al., 2009).

## Acknowledgements

Thanks to: ward managers and the clinical lead for children’s services at Nottingham University Hospitals Trust, healthcare workers, children and their families.

The Authors declares that there is no conflict of interest.

## Supplementary Material

**Supplementary Figure A.**
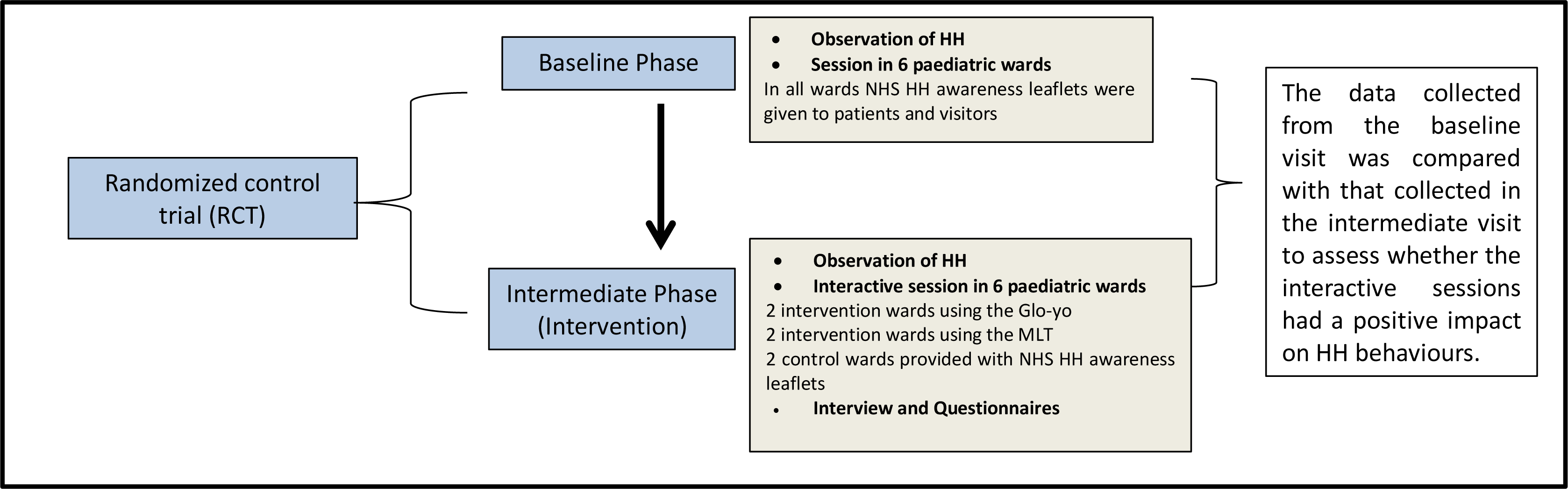
An outline of the study Supplementary Figure B. Glo-yo interactive educational toy.

**Supplementary Figure B.**
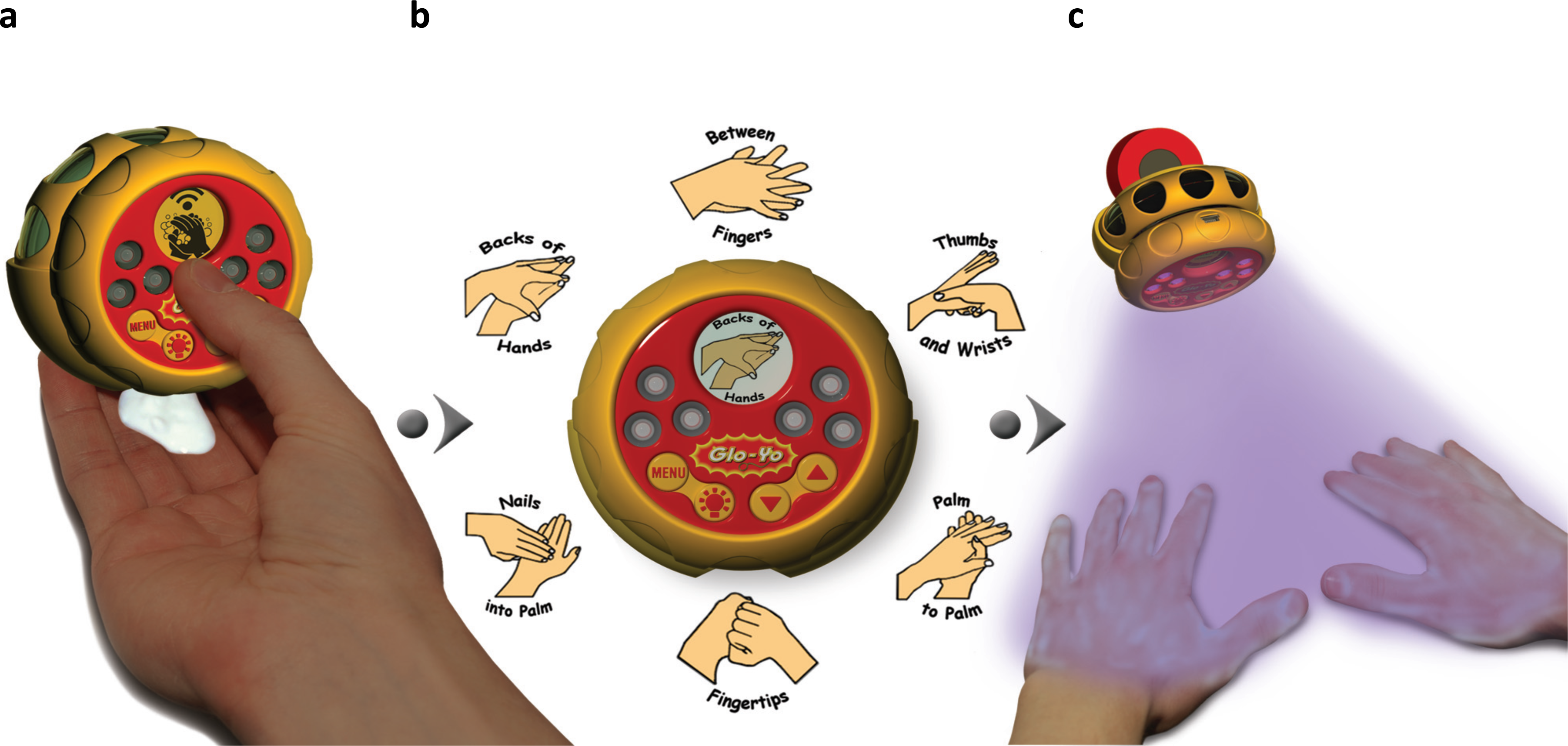
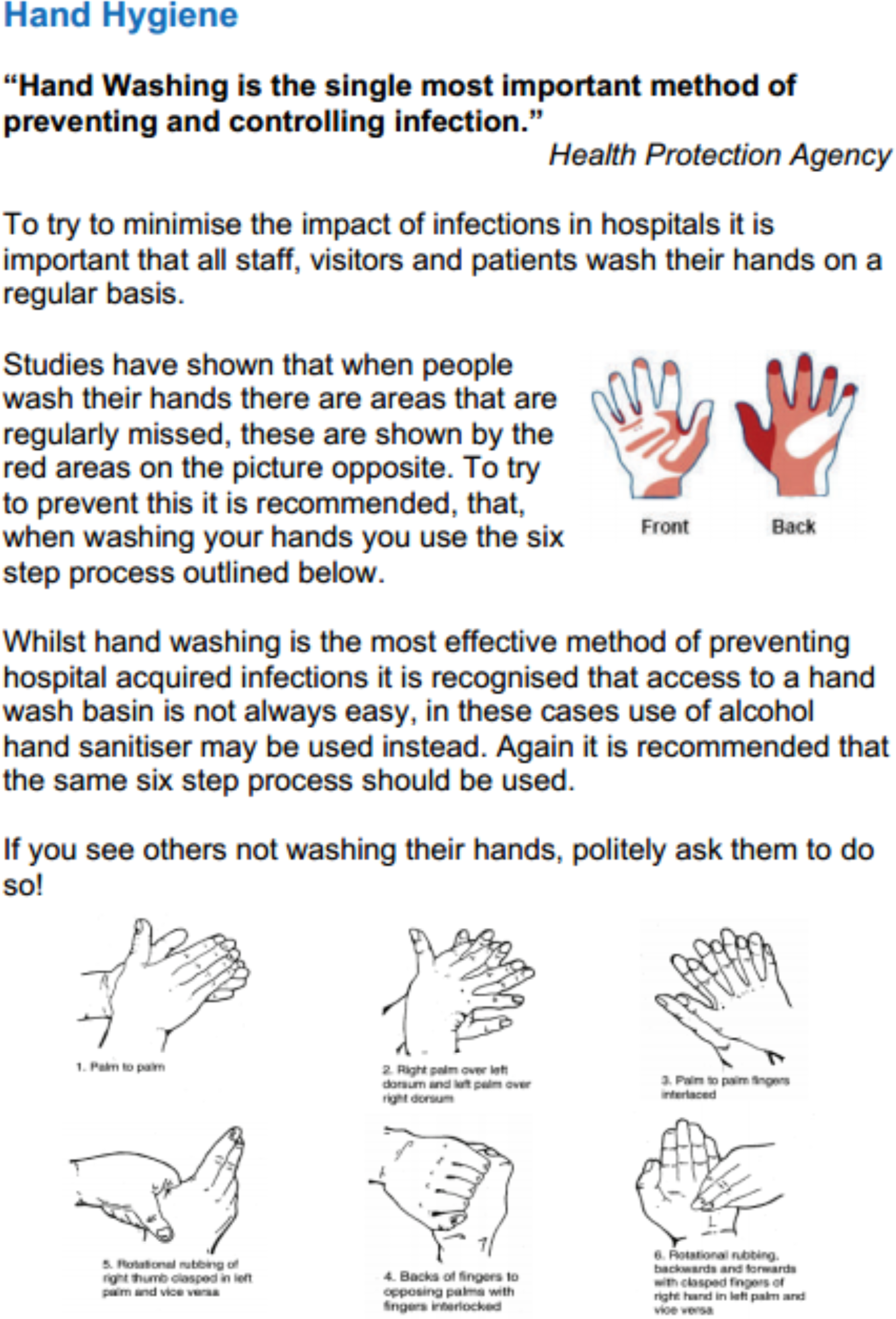
Glo-yo interactive educational toy. (**a**) handheld Glo-yo, (**b)** 6 images of the HHC steps displayed on the screen during 20 seconds, (**c)** UV lights illuminate the iridescent cream on hands as a way to assess the effectiveness of HHC [14].

**Supplementary Figure C.** The leaflet distributed in the control group

**Supplementary Table A.** Description of the Glo-yo, MLT and leaflet used for the RCT. Comparative description of the training aids used in the intervention phase of the trial.

|  | Description | Reference/Link |
| --- | --- | --- |
| **Glo-yo** | The Glo-yo is an interactive toy that helps children learn how to wash their hands better. The toy was designed by an undergraduate engineering student, with the help of school children. The device has a UV light and LDS screen which presents the optimal handwashing regime, timed over 20 seconds as the WHO recommendation for HHC. | http://glo-yo.co.uk/ [14] |
| **MLT** | A video aimed to patients, visitors and HCWs presents the importance of hand hygiene, and the appropriate steps that can reduce the spread of HCAI between patients, visitors and HCWs. The video covers the following: Preventing the spread of germs, when you should clean your hands and hand washing techniques. It was shown to the people on a mobile phone. | The video was produced for the use in hospital and healthcare centres in Nottingham University NHS Trust. Content authors Jacqueline Randle, Heather Wharrad and Richard Windle of the University of Nottingham and Natalie Vaughn of the NHS. Technical development was by Peter Cook (Video) and Fred (Web).<br>http://www.nottingham.ac.uk/nmp/sonet/rlos/placs/cleanyourhands/ |
| **HH Leaflet** | A patient care leaflet that presents the important steps of HH to prevent and control infection. | Supplementary Figure B |

